# PrismExp: Predicting Human Gene Function by Partitioning Massive RNA-seq Co-expression Data

**DOI:** 10.1101/2021.01.20.427528

**Authors:** Alexander Lachmann, Kaeli Rizzo, Alon Bartal, Minji Jeon, Daniel J. B. Clarke, Avi Ma’ayan

## Abstract

Gene co-expression correlations from mRNA-sequencing (RNA-seq) can be used to predict gene function based on the covariance structure that exists within such data. In the past, we showed that RNA-seq co-expression data is highly predictive of gene function and protein-protein interactions. We demonstrated that the performance of such predictions is dependent on the source of the gene expression data. Furthermore, since genes function in different cellular contexts, predictions derived from tissue-specific gene co-expression data outperform predictions derived from cross-tissue gene co-expression data. However, the identification of the optimal tissue type to maximize gene function predictions for all mammalian genes is not trivial. Here we introduce and validate an approach we term Partitioning RNA-seq data Into Segments for Massive co-EXpression-based gene function Predictions (PrismExp), for improved gene function prediction based on RNA-seq co-expression data. With coexpression data from ARCHS4, we apply PrismExp to predict a wide variety of gene functions, including pathway membership, phenotypic associations, and protein-protein interactions. PrismExp outperforms the cross-tissue co-expression correlation matrix approach on all tested domains. Hence, PrismExp can enhance machine learning methods that utilize RNA-seq coexpression correlations to impute knowledge about understudied genes and proteins.

## Introduction

The automated annotation of gene function can be derived by combining prior knowledge manual annotations with mRNA co-expression patterns based on the concept of guilt-by-association. In general, it is well established that genes that co-express display functional relatedness (1–3). Proteins with high mRNA expression correlations, or high mutual information (4), are often found in the same macro-molecular complexes, cell signaling and metabolic pathways (5, 6). Since proteins predominantly function through the formation of protein complexes and regulatory networks, their activity depends on their expression in different cellular and tissue contexts. Tight control of protein complex stoichiometry is needed for their correct formation (7). While all cells and tissues carry out general housekeeping processes, cells and tissues also have to fulfill specialized roles and subsequently differ in their pathway and biological function composition. Due to their cell and tissue specificity, considering gene regulatory networks in the right cellular and tissue context is important (8). Reverse engineering of regulatory mechanisms with tissue agnostic approaches lacks fidelity to capture nuanced gene-gene and protein-protein interactions accurately. With the availability of RNA-seq data from resources such as ARCHS4 (9), Genotype-Tissue Expression (GTEx) (10), or The Cancer Genome Atlas (TCGA) (11) featuring cell type and tissue specific gene expression information, it is feasible to leverage the context specific gene expression patterns for improved gene function predictions.

Over decades of biochemistry, pharmacology, molecular and cell biology research, many mammalian genes have been thoroughly characterized. However, there are still many genes that remain obscured due to a lack of research interest. Extensive gene function annotations can be accessed from resources such as the Gene Ontology (12), Human Phenotype Ontology (13), UniProt (14), Harmonizome (15) and Enrichr (16). The Gene Matrix Transposed (GMT) file format, used by Gene Set Enrichment Analysis (GSEA) tools (17) provides a convenient way to organize knowledge about genefunction associations. Each row in a GMT file corresponds to a biological function, followed by a set of associated genes. The assumption is that these gene-sets are generally incomplete, reflecting our partial knowledge about the involvement of genes and their protein products in biological processes.

Computational methods that can utilize gene function annotations to impute the function of understudied genes are an efficient way to form novel hypotheses that can lead to new discoveries. High-content genome-wide data, such as RNA-seq, includes information about under-studied genes and is not biased towards well-studied genes. Combining prior knowledge gene annotations with high-content genome-wide data can translate existing knowledge about well-studied genes to the under-studied genes. Significant efforts are made to produce data and develop tools to shed light on these understudied genes (18–20). One such effort is the NIH Common Fund Program Illuminating the Druggable Genome (IDG) (21). IDG focuses on understudied genes from three prominent gene families: protein kinases, G-protein coupled receptors (GPCRs), and ion channels. These families contain the most promising druggable protein targets with many understudied members (22).

In the past, we have developed methods to predict gene function for all human genes, including members of these three families, using co-expression data that is tissue agnostic. It is reasonable to expect that by considering the tissue context of co-expression data the quality of such predictions will improve. However, the utilization of tissue specific gene expression patterns poses practical limitations:

I. When making predictions about a specific biological function, the correct context of the particular function has to be considered. With potentially hundreds of relevant tissues to choose from, tissue specific predictions require significant domain knowledge.
II. While some gene expression resources are well organized into individual tissues, for example, GTEx (10) or TCGA (23), these resources only cover a fraction of all human tissues and cell types. On the other hand, more diverse datasets such as ARCHS4 (9) lack accurate tissue classification of individual samples.
III. Gene function prediction algorithms that utilize coexpression data are applied to predict thousands of diverse biological functions. Thus, manual input for individual cases is not feasible.

To address the above limitations, while still leveraging the advantage of tissue specific gene function prediction using co-expression patterns, we developed a novel machine learning approach called Partitioning RNA-seq data Into Segments for Massive co-EXpression-based gene function Predictions (PrismExp). PrismExp generates a high dimensional feature space from large unlabeled RNA-seq mRNA gene expression data. The generated feature space automatically encodes tissue specific information via vertical partitioning of the data matrix. We demonstrate that this approach significantly improves gene function predictions when compared to methods that do not partition the data matrix.

## Materials and Methods

### A. Data resources

To compute pairwise gene-gene mRNA co-expression correlations, we used the mouse ARCHS4 RNA-seq gene expression dataset. We chose mouse over human because benchmarking (9) demonstrated that predictions made with the mouse co-expression data slightly outperformed predictions derived from human co-expression data. The ARCHS4 mouse dataset is comprised of 284,907 uniformly aligned RNA-seq gene expression samples publicly available from GEO (24). For benchmarking, we utilized all 163 available gene-set libraries from Enrichr (16). For more detailed analysis, we used a subset of the gene-set libraries from Enrichr (Table 1). In total, the collection of the genesets in these five libraries contain 19,906 gene functions.

**Table 1.** Gene-set libraries from Enrichr with the number of gene-sets and unique genes. Literature based gene-set libraries are manually curated and non-literature based libraries are derived from directly from experimental outcomes.

| GMT |  | Description | Literature | Sets | Genes |
| --- | --- | --- | --- | --- | --- |
| ChEA 2016 | (25) | Transcription factor targets from ChIP-seq experiments | ✗ | 645 | 17022 |
| KEA 2015 | (26) | Kinase-substrate interactions | ✗ | 428 | 2904 |
| MGI phenotype | (27) | Gene-phenotype associations from phenotype screens | ✗ | 5261 | 10390 |
| hu.MAP | (28) | Protein-protein interactions | ✓ | 995 | 2117 |
| GO: Biological Process | (12) | Manually curated ontology of gene function | ✓ | 5103 | 12096 |
| GWAS Catalog 2019 | (29) | Manually curated SNP-trait associations from literature | ✓ | 1737 | 10744 |

### B. Generating the gene-gene mRNA co-expression matrices

We first filter out genes with low gene expression, resulting in *G*, a set of genes with expression values across samples. The threshold to include genes in *G* requires a gene to have at least 20 reads in 1% of all samples. This filtering step is intended to remove pseudo-genes and genes with very low levels of transcript abundance. PrismExp automatically identifies a set of *N* clusters defined as *C* using a k-means clustering algorithm. For each cluster defined by the k-means clustering *C*, we apply quantile normalization on the log2 transformed gene counts to correct for library size variability. For each resulting normalized gene expression matrix, we calculate the pairwise gene-gene correlations for genes in *G*, *C_i_* for cluster *i* using Pearson correlation. Additionally, we calculate the Pearson correlation coefficients between genes across a random selection of 10,000 samples, *C_global_*. The set of correlation coefficient matrices is: *C* = {*C*_0_, *C*_1_,…, *C_k_, C_global_*}.

### C. Computing gene × gene-set features

Gene × geneset features are scores for the strength of association between a gene and a gene-set pair. For a gene *g_x_* ∈ {*g*_1_,…, *g_L_*} and a gene-set *p_y_* ∈ {*p*_1_,…, *p_M_*} the algorithm generates a set of features *S_k_* used in the final prediction step. A gene-set library *S* ∈ *L* × *M* is defined as a binary matrix with genes as rows and gene-sets as columns.

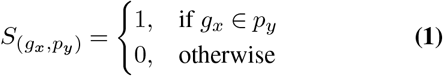

Given *N* gene co-expression matrices *C_k_* ∈ {*C*_1_,…, *C_N_*} and a gene-set library *S*, the corresponding gene-function association features 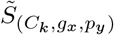 are defined as,

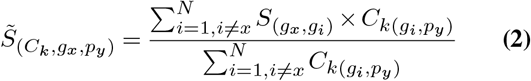

Alternatively, a gene-set library can also be defined as a nonbinary adjacency matrix. Given *N* co-expression matrices, each pair of gene and gene-set/function has a set of features,

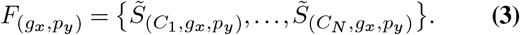

### D. Training a Random Forest regression model

PrismExp takes *F_gx,py_* as input features to classify whether *g_x_* should be annotated with *p_y_* as a functional association. Support Vector Machine (SVM), artificial neural networks (ANN), and other machine learning algorithms are suitable to apply for constructing predictive models in this context. PrismExp utilizes a Random Forest regression model (30) to accomplish this task. We also tested several other machine learning models (Support Vector Machine, Multi-layer Perceptron regressor, logistic regression, XGBoost, Gaussian Naive Bayes, k-Nearest Neighbor, and Linear Discriminant Analysis) (31–37) and all performed comparable. The model is trained with the following data,

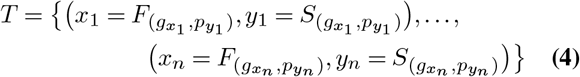

### E. Python package and web interface

PrismExp is available as a Python package. The package supports programmatic prediction of gene function from gene expression data utilizing the data from ARCHS4 (9) stored in HDF5 format (38). Predictions can be made for any gene-set library. The PrismExp package also includes gene-set libraries to use directly from the collection of libraries available from Enrichr (16). The PrismExp Python package offers parameter customization, for example, the user can adjust the number of gene expression clusters *C*. In addition to the PrismExp Python package, PrismExp is also available as an Appyter (https://appyters.maayanlab.cloud/PrismEXP/). Appyters convert Jupyter Notebooks to web-based bioinformatics applications. The PrismExp Appyter enables users to execute the PrismExp computational pipeline using their own data in the cloud without coding. The PrismExp Appyter has access to 51 pre-computed correlation matrices stored in Amazon Web Services (AWS) S3. The Appyter implementation of PrismExp directly extracts the required correlation data from the pre-computed matrices. A user interface enables the selection of a gene of interest, as well as the selection of any gene-set library for target predictions. Alternatively, the user can upload their own custom gene-set library in GMT format.

**Fig. 1.**
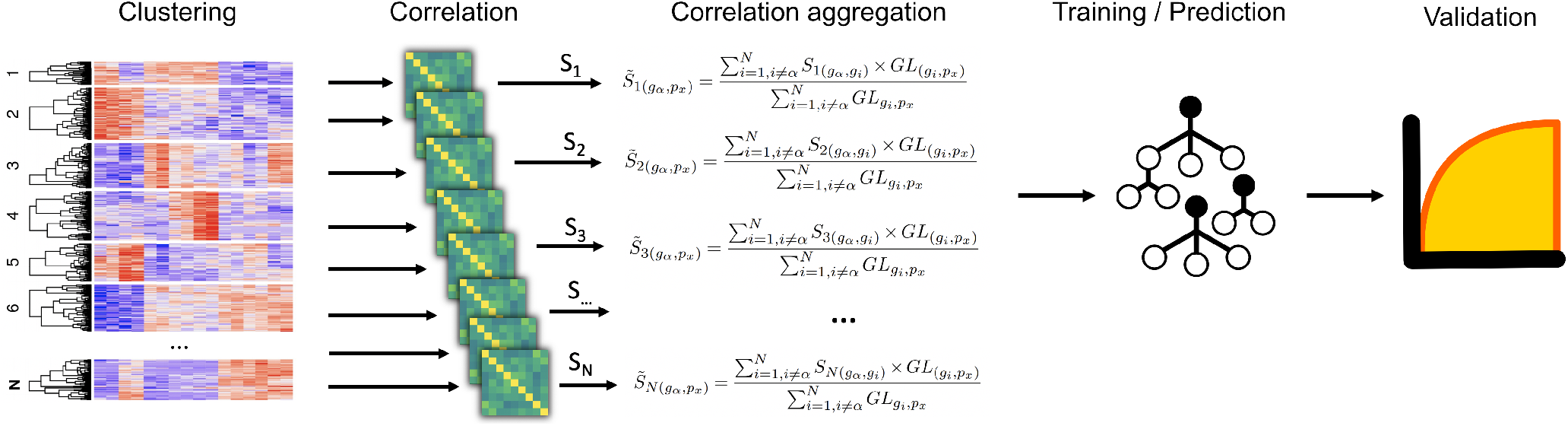
PrismExp workflow for gene function prediction.

### F. Validation

To assess the quality of the predictions made by PrismExp, we asked how well the prediction method can retrieve the known biological functions of a gene. The PrismExp prediction output is a matrix with genes as rows, and biological functions as columns. The higher a value in the matrix, the more likely the corresponding gene is associated with the respective biological function. Thus, there are two ways to assess the quality of the predictions made by PrismExp:

- Rank the values of each row in descending order
- Rank the values of each column in descending order

Ranking the values for each row is equivalent of ranking gene functions for each gene from likeliest to least likely. Since there is prior knowledge about the associations of genes with biological functions, these incidences are marked as positives. These positives are expected to be highly ranked. Biological functions that were not previously associated with a gene are considered false, and are expected to have lower ranks. PrismExp calculates an area under the receiver operating characteristic (AUROC) curve for each gene. These AUROC values are used to compare the quality of the predictions by different methods, model parameters, and underlying data sets.

Alternatively, we can rank the genes for each biological function by the most likely to the least likely. Similarly, we can calculate the AUROCs by labeling the previously known associations. Both sets of AUROC scores, ranking gene for biological function and ranking biological functions for genes, are critical to consider when evaluating and validating various sources of data and model parameter settings.

## Results

### G. Gene function prediction workflow

PrismExp produces predictions about gene-function associations at the genome-wide scale. The predictions are based on prior knowledge about gene-function associations and gene-gene co-expression data. The PrismExp workflow is made of two main branches: initialization and prediction 1. During initialization, PrismExp generates the required correlation matrices and pre-trains a Random Forest regression model. The prediction step takes as input a gene-set library stored in Gene Matrix Transposed (GMT) format, and uses the precomputed correlation matrices and regression model to make predictions. To make a prediction, PrismExp calculated the average correlation of a gene with a gene-set for each correlation matrix. The resulting feature vector is used as the input for the Random Forest model. As an optional step, PrismExp can validate the prediction performance for a given biological function, or for a specific gene using AUROCs.

### H. Performance and hardware requirements

PrismExp is optimized to run on standard desktop computers utilizing multiple cores for the clustering and correlation calculations. Since PrismExp is using large matrices, memory consumption should be considered. For a realistic use-case, using the ARCHS4 mouse gene co-expression dataset, the memory requirements are 8GB. PrismExp was benchmarked using an Intel i7-8750H with 6 cores and 32GB of RAM using Python 3.8.5. PrismExp was also tested on an AWS r5ad.2xlarge instance with 8 cores and 64GB of RAM running Ubuntu. Both tests produce comparable performance (Figure S1). The memory consumption and compute time from the five different stages of PrismExp execution, training it with 50 gene expression clusters, shows that gene filtering and sample clustering is completed in 50 minutes. Calculating the correlation matrices takes the longest time, 122 minutes or 2.4 minutes per cluster. Reducing or increasing the number of clusters affects run-time linearly based on the number of clusters. The prediction step depends on the number of genes, the number of gene-sets, and the average gene-set size of the gene-set library. This is in addition to the number of gene expression clusters. This is determined based on prediction and training applied using the Gene Ontology Biological Processes 2018 library from the Enrichr gene-set library catalog. This gene-set library contains 5,103 gene-sets, 14,433 genes, and has an average of 36 genes per set. Training the Random Forest model with 40,000 positive samples and 200,000 negative samples take about 50 minutes to complete. The steps shown above have to be performed only once to initialize the prerequisite data for making multiple predictions. Custom gene-set libraries can be used to train the Random Forest regression model. The only step that needs to be performed each time for each gene-set library is the prediction step, which calculates the average correlation of a gene to all gene-sets in the library for each correlation cluster.

The number of gene expression clusters, and the resulting coexpression correlation matrices, impact the final prediction quality. We compared the prediction quality for 5, 10, 25, 50, 100, and 300 clusters (Figure 2). These results are compared to predictions made with the global correlation matrix that is not partitioned into clusters. Overall, we observe a significant improvement in the average AUROCs for using the partitioning method over the global co-expression correlation matrix. While the performance of the global correlation matrix outperforms PrismExp for low numbers of clusters, PrismExp significantly beats the prediction quality made by the global correlation matrix when more clusters are considered. We applied a paired sample Student t-test on the average AUROCs from 163 gene-set libraries to compare the performance of a single correlation matrix to applying PrismExp with 301 clusters. The results confirm significant improvement for using the partitioning approach, *p* = 1.401*e*^−37^ and *p* = 2.397*e* – 31 for sets and genes respectively. The increase in average AUROCs achieved by a higher cluster count can be represented roughly by a logarithmic function of the shape *f*(*x*) = *a* × *log*_2_(*bx*) + *c* 2. This fitting suggests that the number of clusters is logarithmically proportional to the average AUROCs. Increasing the number of clusters beyond 300 only marginally improves the predictions while increasing the computation time considerably.

**Fig. 2.**
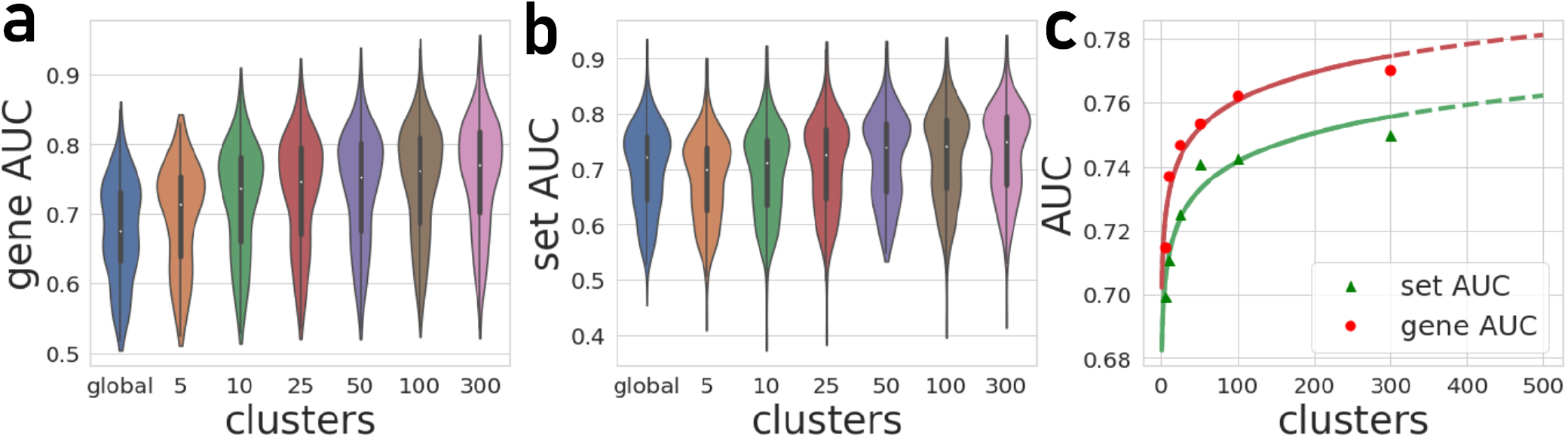
**a)** Average AUROCs of gene-sets for 163 gene-set libraries. **b)** Average AUROC of gene-sets and predicted AUROC performance by the number of gene expression clusters. **c)** Average AUROCs of genes for the 163 gene-set libraries. **d)** Average AUROC of gene-sets and predicted AUROC performance by co-expression clusters.

By projecting the high dimensional feature-space of PrismEXP using t-SNE (39) we observe distinct clusters (Figure 3). In these plots, each point represents a gene x gene-set pair. Gene × gene-set pairs with prior known associations are organized into a gradient of average correlation from low on the left to high on the right (Figure 3 **b**). A similar gradient from left to right is observed for the PrismExp predictions, following the true positives sample distribution (Figure 3 **c**). The features derived from the 301 co-expression correlation matrices, in combination with the Gene Ontology gene-set library as the training set, show that individual gene co-expression matrices contribute to the performance of the PrismExp model (Figure 4 **a** and **b**). The feature importance of each co-expression correlation matrix is related to the prediction performance when using the average correlation of the correlation matrix alone. Co-expression correlation matrices that contribute a high level of feature importance also perform better individually than matrices with low feature importance contribution. The global co-expression correlation matrix computed from 10,000 randomly selected samples extracted from the ARCHS4 resource has the highest single matrix performance and the highest feature importance contribution.

**Fig. 3.**
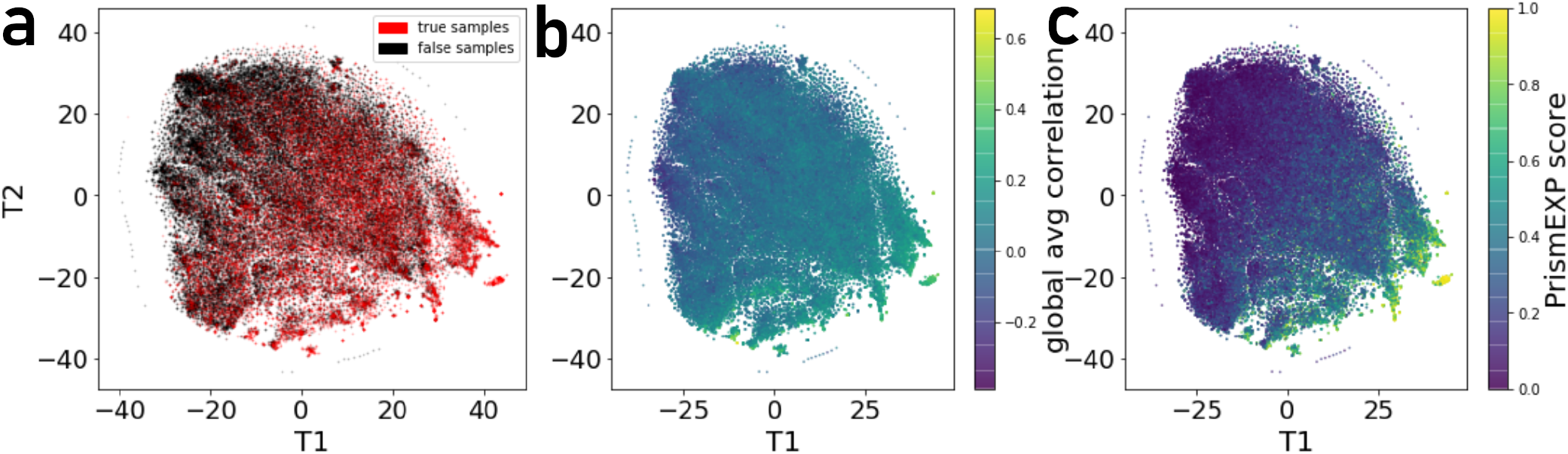
t-SNE visualization of 301 dimensional feature-space of 17,000 gene x gene-set pairs. **a)** Distribution of gene x gene-set pairs with known association (true positives). **b)** Average correlation scores of gene x gene-set pairs relative to the high dimensional feature space. **c)** PrismExp scores for gene x gene-set pairs relative to the high dimensional feature space.

**Fig. 4.**
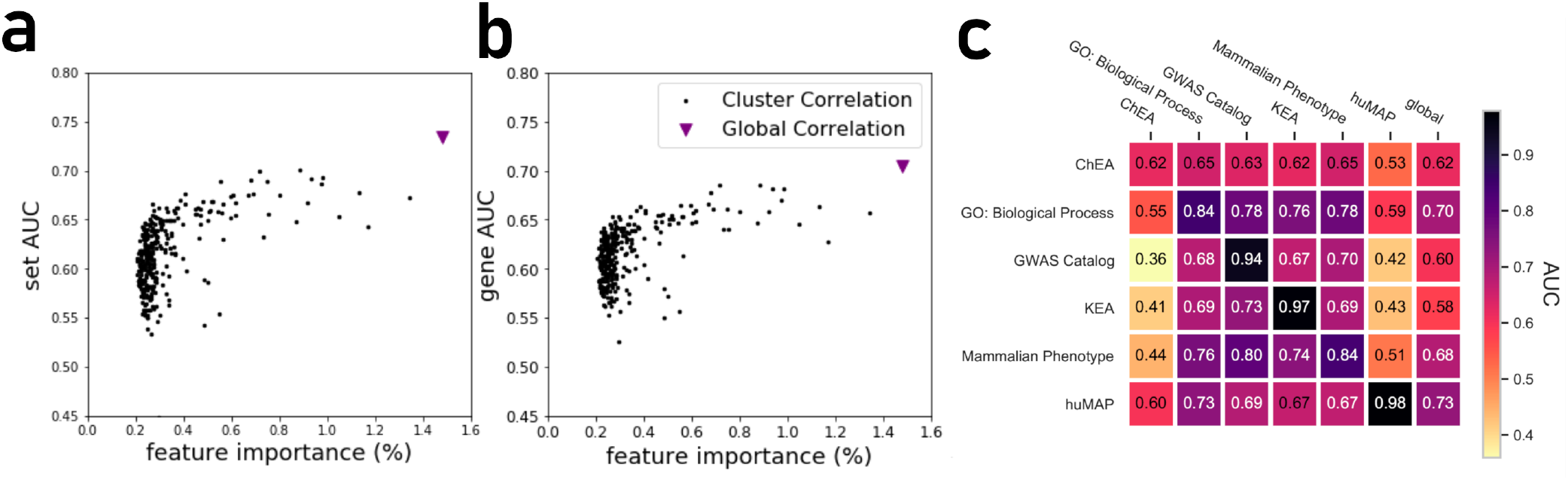
**a)** Feature importance of individual correlation matrices relative to their prediction performance of ranking genes based on predicted known associations with biological functions. **b)** Feature importance of individual correlation matrices relative to their prediction performance in ranking gene-sets based on their associations with genes. **c)** PrismExp Random Forest model prediction performance evaluated as the average gene AUROC when trained on six gene-set libraries. The plot also includes prediction performance when using the global single correlation matrix.

It is expected that a model trained with a specific gene-set library will produce high prediction performance when predicting gene functions from that same library. However, we next asked how well a model trained with one gene library can predict terms from another library (Figure 4 **c**). A heatmap that compares average AUROCs illustrates that the six PrismExp models trained on one gene-set library listed in 1 can be used to predict terms in other libraries. The prediction performance of a model trained on the same geneset library is always the best, as expected. For example, a model trained on the hu.MAP gene-set library achieves an average gene AUROC of 0.98 when applied to make predictions for the same gene-set library. However, this model does not achieve good predictions for any of the other tested geneset library. The results suggest that the hu.MAP gene-set library contains different co-expression patterns than the other libraries, and thus do not generalize gene function predictions across a diverse set of libraries. However, the model trained with GO Biological Processes (BP) achieves good prediction performance when used to predict other libraries’ functions. The GO-BP model also outperforms the single matrix predictions using all tested gene-set libraries, suggesting that the GO-BP model is generalizable across multiple gene-set libraries S2.

### I. Predicting under-studied druggable targets with PrismExp

The Illuminating the Druggable Genome (IDG) project is focused on furthering our knowledge about understudied potential protein targets from the ion channel, kinase, and GPCR gene families (40). Here we applied PrismEXP to predict the gene functions for one understudied member from each of these three drug-target-rich families. The most significant alpha-kinase 3 (ALPK3) associations are illustrated in a network diagram that connects ALPK3 with the top predicted GO terms and other relevant genes (Figure 5a). To date, there are only 20 publications that mention ALPK3. One of these, reported that biallelic loss of function of this gene is implicated in earlyonset cardiomyopathy (41). The top gene functions predicted for ALPK3 by PrismExp are skeletal muscle contraction (p=1.244e-15), myosin filament assembly (p=2.8701e-14), cardiac muscle hypertrophy (p=3.526494e-13), cardiac muscle fiber development (p=8.0835e-13), and striated muscle contraction (p=1.0353e-12). In addition, ALPK3 is highly related to well-studied genes that form the sarcomere, namely TTN, MYOM1, MYOM2, and MYOM3. Hence, the understudied kinase ALPK3 has an important role in cardiac function that is highly under-appreciated. For GPR183, PrismExp predicts involvement in immune system-related processes. The top 5 terms are B-Cell activation (p=9.5052e-15), lymphocyte chemotaxis (p=3.8298e-09), positive regulation of B-Cell proliferation (p=4.683e-09), T-Cell differentiation (p=5.7275e-09), and T-Cell activation (p=1.1794e-08). GPR183 highly associated genes are the immune-response cytokines IL2, IL4, and CCL19 (Figure 5b). The predicted associated kinases for GPR183 are the non-receptor tyrosine kinases LCK, TXK, BLK, and IT; suggesting a direct or an indirect regulatory role for tyrosine phosphorylation to regulate the function of this GPCR. Finally, for the ion channel CLIC5, PrismExp predicts roles in inner ear receptor stereocilium organization (p=3.285e-20), sensory perception of mechanical stimulus (p=1.7546e-07), regulation of systemic arterial blood pressure (p=1.3696e-06), vasoconstriction (p=1.1852e-05), and positive regulation of vasculogenesis (p=2.4557e-05) (Figure 5c). Loss of CLIC5 has been associated with deafness (42). Hence, PrismExp is highly predictive for illuminating knowledge about under-studied genes and proteins.

**Fig. 5.**
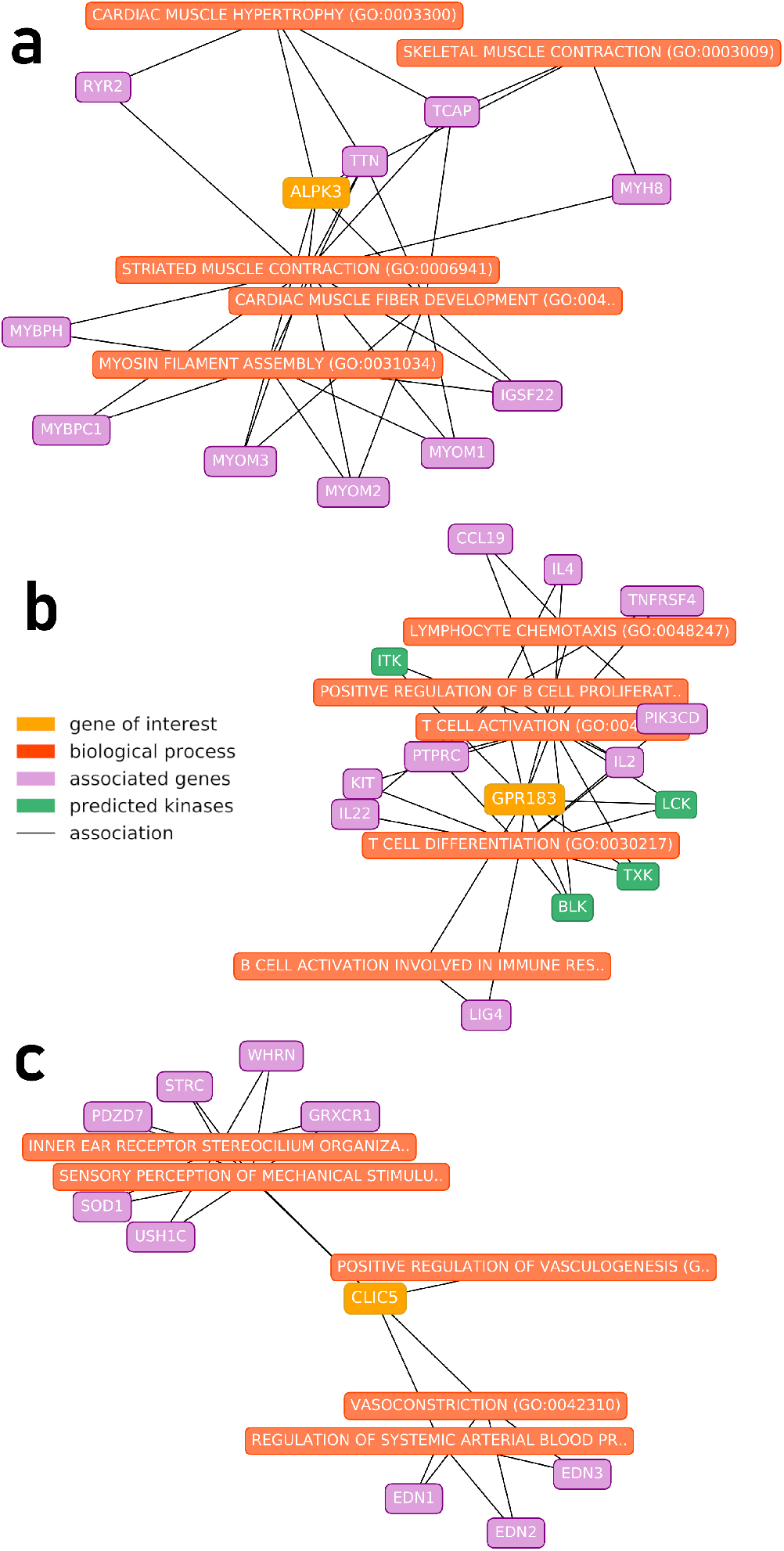
Top 5 predicted Gene Ontology Biological Processes (orange) and most associated other genes (purple) for understudied targets (yellow) **a)** The protein kinase ALPK3, **b)** The GPCR GPR183, and **c)** the ion channel CLIC5. Green nodes represent predicted kinases for GPR183.

## Summary

Here we demonstrated that we could improve the inference of gene function by automatically partitioning the ARCHS4 gene-gene co-expression correlation matrix. Such automated partitioning is likely positively correlated with tissue and cell type partitioning but without the need for metadata labeling. The method treats each gene expression cluster as a source for features to construct machine learning models for gene function prediction. This approach deviates from prior work that required correct labeling of the gene expression profiles, for example, by text mining the metadata (43). Another advantage of PrismExp is that the RNA-seq data is available for all genes. Thus, it is suitable for predicting biological functions for understudied genes, including non-coding genes. The predictions performed by PrismEXP can be used for hypothesis generation and guidance in the design of wetbench experiments. Specifically, PrismExp generates ranks for genes associated with biological functions, or ranks of biological functions for genes, facilitating many use cases. If a gene of interest is already associated with a biological function, a phenotype, or a disease, PrismExp can be used to highlight other genes with similar profile.

As with all machine learning applications, there is a risk of over-fitting the PrismExp model to patterns embedded within one specific gene-set library used for training. Hence, we aimed to evaluate how generalizable the models created using one library are to make predictions of known gene functions from other libraries. We confirmed that the models are, in general, generalizable. In addition, we observed a significant improvement in the predictions for hu.MAP and KEA due to the partitioning the samples for these libraries. Further research should be able to discern why such significant improvement is observed for these datasets. One postulation is that protein-protein interactions and kinase-substrate phosphorylations are context specific.

In this study, all predictions were made based on features that are derived from RNA-seq mRNA co-expression data. However, other omics data modalities could be incorporated into the same framework. By adding other sources of gene-gene similarity, additional features can be added to the model. We have explored the use of other gene similarity matrices for gene function prediction for other applications. For example, for developing Geneshot (18), we predicted gene functions with features from gene-gene co-mentions in the literature, and co-occurrence in user-uploaded gene-sets to Enrichr (16). However, we never attempted to partition the gene-gene similarity matrices from these resources.

Finally, to make the PrismExp accessible to the community, we developed a Python package that enables computational biologists access the PrismExp method. In addition, we also provide access to the predictions that are made by PrismExp via an Appyter. The Appyter provides access to gene function predictions for users without programming skills. In the future, we plan to add the PrismExp predictions to the ARCHS4 (9) gene pages.

## ACKNOWLEDGEMENTS

This work was partially supported by the National Institutes of Health (NIH) grants U54-HL127624 (LINCS-DCIC) and U24-CA224260 (IDG-KMC) to AM.

## Supplementary Note 1: Supplementary Material

**Fig. S1.**
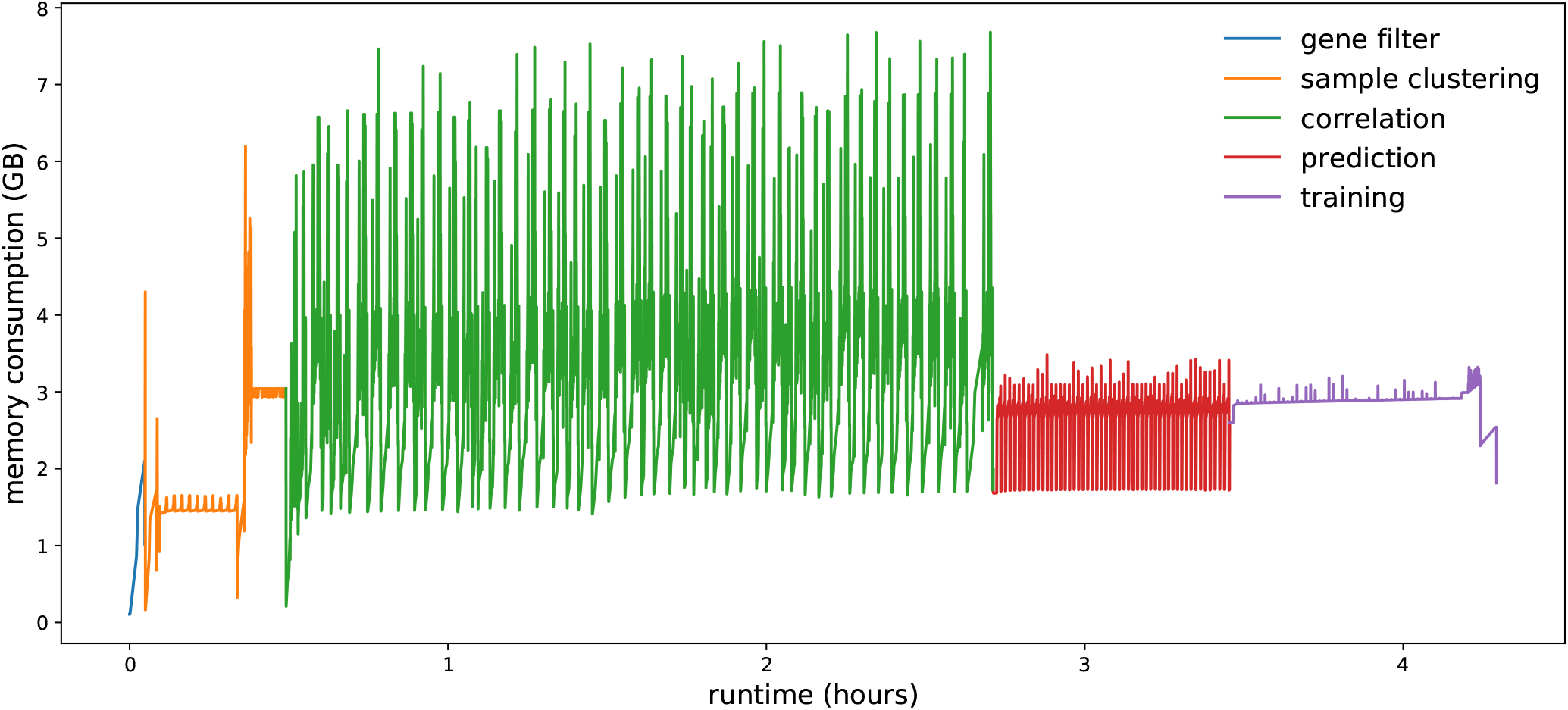
PrismExp runtime and memory consumption for 50 gene expression clusters.

**Fig. S2.**
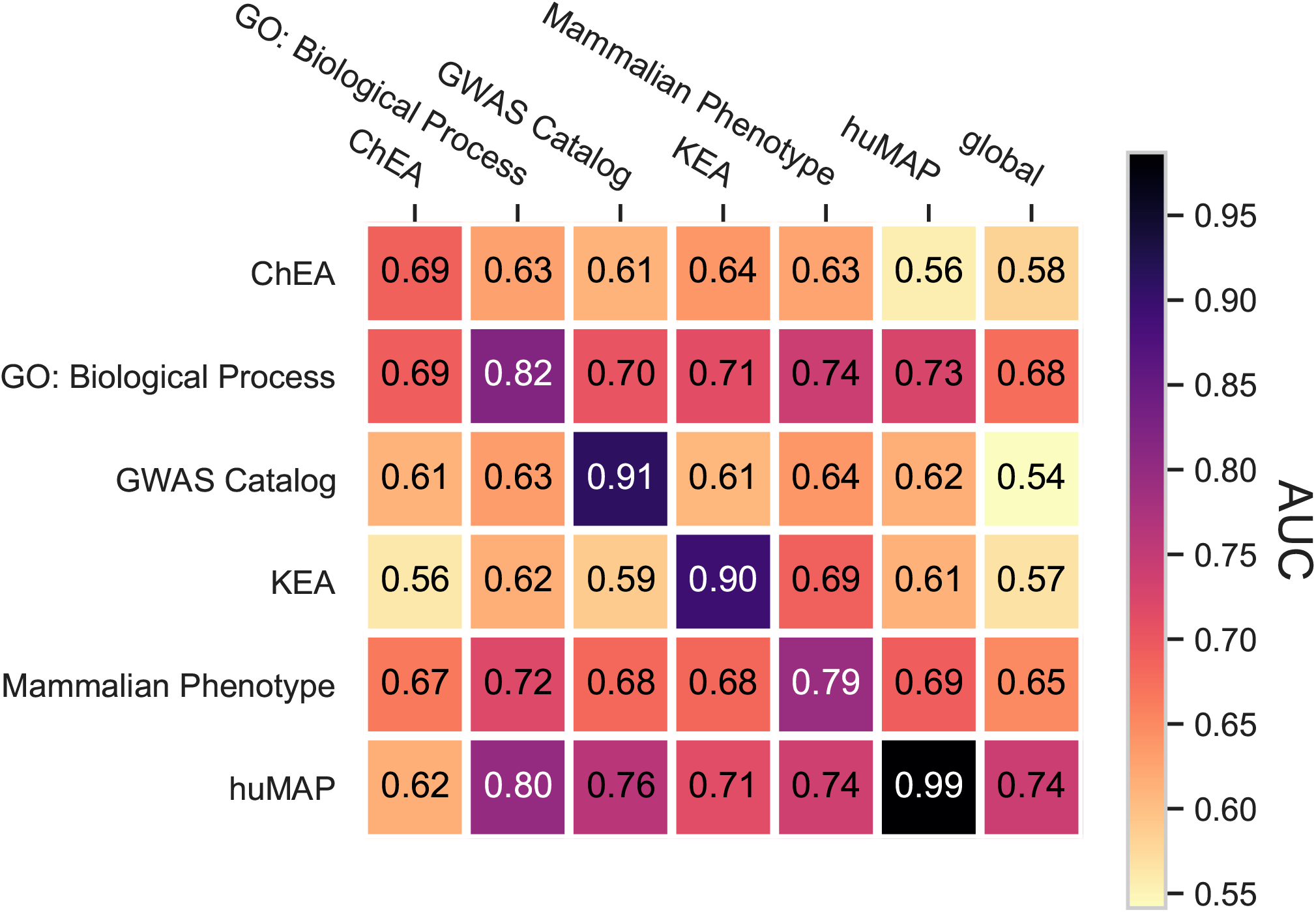
Random Forest model prediction performance (average set AUROC), when PrismExp is trained on six gene-set libraries as well as prediction performance of single correlation matrix (global).

